# Developmental retrotransposon activation primes host immunity for future viral-clearance

**DOI:** 10.1101/2020.08.23.263293

**Authors:** Lu Wang, Lauren Tracy, ZZ Zhao Zhang

## Abstract

Transposons are thought to be largely suppressed under physiological conditions, ensuring that their mobilization is a rare event. By tracking mobilization, we show that during metamorphosis at the *Drosophila* pupal stage, the *Gypsy* retrotransposon selectively mobilizes in regenerating tissues. In the newly formed tissues, this wave of *Gypsy* activation primes the host’s innate immune system by inducing the production of <u>a</u>nti<u>m</u>icrobial <u>p</u>eptides (AMPs). Moreover, early immune-priming functions of *Gypsy* are essential for combating viral invasion in adult flies: flies with *Gypsy* being silenced at the pupal stage are unable to clear viruses and succumb to viral infection. Our data reveal that regulated activation of transposons during animal developmental endows a long-term benefit in pathogen warfare.

---

Retrotransposons comprise 38% of the human genome. Although recent studies suggest positive effects of retrotransposon activation, such as contribution to neuronal genomic mosaicism or as genomic regulatory sequences (1–5), mobilization of these elements generates DNA damage and mutations, causing diseases and potentially driving aging (5–7). Assuming that they are largely suppressed, extensive efforts led to the identification of mechanisms that silence retrotransposons in both germline and somatic cells in the past (8, 9). Here we seek to explore, even under strict regulation, what the retrotransposon activity is at the organismal level.

Encouraged by our previous work on monitoring mobilization during oogenesis (10, 11), we sought to systematically characterize retrotransposon jumping events in somatic tissues of *Drosophila*, a powerful genetic model with compact genome and well-curated transposon annotation. To achieve this, we chose to explore 9 retrotransposon families with the capacity to mobilize: *3S18*, *412*, *Blood*, *Burdock*, *Copia*, *Doc*, *I-element*, *Mdg1*, and *Gypsy* (10, 12–19). For each family, we engineered a corresponding eGFP mobilization reporter (Fig. 1A and Fig. S1A). Similar to the reporters designed in previous studies (10, 20, 21), an eGFP cassette was inserted into the retrotransposon in an antisense direction. This cassette contains a disruptive intron that can be spliced during transcription of the retrotransposon, but not of eGFP. As such, the reporter can only express eGFP after intron-spliced transposon mRNAs are used as templates to make new copies of DNA (Fig. 1A and Fig. S1A). To minimize the positional effects from chromatin on transposon activity, we generated fly alleles that carry each eGFP reporter at 2 different genomic loci.

**Fig. 1.**
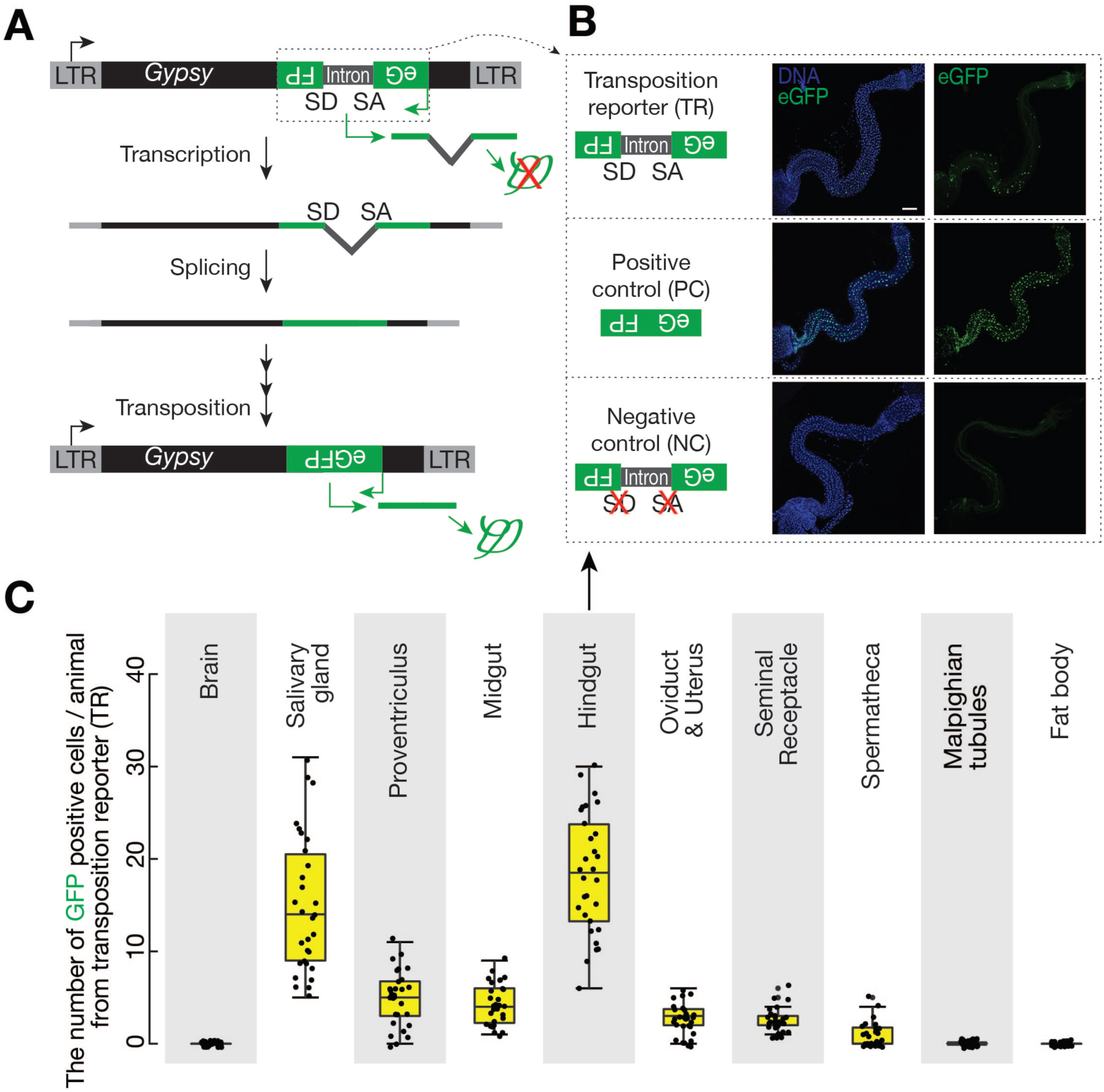
Monitoring *Gypsy* mobilization in somatic cells via a transposition reporter. **(A)** Schematic design of an eGFP reporter to monitor *Gypsy* mobilization. The eGFP reporter is inserted in the non-coding sequence of *Gypsy* in the anti-sense orientation. eGFP is disrupted by an intron, which is in the same direction as *Gypsy* but opposite direction as eGFP. The intron cannot be spliced during eGFP transcription. *Gypsy* mobilization, which generates a new copy of the transgene without the disrupting intron, results in eGFP expression. **(B)** *Gypsy* transposition reporter produces eGFP in hindgut cellsin 2-4 dayold adult flies. An intronless construct is used as a positive control to test the potential sensitivity. A construct with mutated splicing acceptor and donor sites serves as a negative control. **(C)** Quantification of the number of eGFP positive cells from *Gypsy* transposition reporter among different tissues from 2-4 day old adult flies.

We next examined 7 somatic tissues from 2-4-day-old flies to examine eGFP signals from the mobilization reporters: brain, salivary gland, proventriculus, midgut (equivalent to human small intestine), hindgut (equivalent to human colon), Malpighian tubule (equivalent to human kidney), and fat body (equivalent to human liver). Among the 9 retrotransposon families, 8 displayed no detectable eGFP signals (Fig. S1B). By contrast, the reporter for *Gypsy* produced eGFP signals in salivary gland, proventriculus, midgut, and hindgut (Fig. 1B, 1C, Fig. S1B, and S1C). Thus, our data indicate that *Gypsy* still can mobilize in somatic tissues. Based on the reporter design, both original cells harboring mobilization events and their progeny can produce eGFP. The mobilization events we detected could occur either in the adult cells we examined, or in their developmental precursors.

As a holometabolous insect, *Drosophila* proceeds through four different life stages: the embryo (~1 day); larva (~4 days); pupa (~4 days); and adult (~2 months). To determine when *Gypsy* mobilizes, we first examined the most highly labelled tissue, hindgut, at these stages (Fig. 1C). We discovered that *Gypsy* selectively becomes active at pupal stage (Fig. 2 and Fig. S2). Notably, metamorphosis is initiated in pupae, during which many tissues from fly larvae are first destroyed and then regenerated to produce adult structures, such as salivary gland and gut (22). For the first 20 hours after puparium formation (APF), the stage of hindgut degeneration, we detected no *Gypsy* mobilizations (Fig. 2A and Fig. S2A). However, at 24 hours APF, we observed eGFP signals from the mobilization reporter in newly formed hindguts (Fig. 2A). As the hindgut grew during metamorphosis, we detected more cells harboring mobilization events (Fig. 2A). Similar to hindgut, *Gypsy* does not appear to mobilize in the degenerating salivary gland, proventriculus, and midgut at early pupal stage, but mobilizes in these tissues during their developmental regeneration (Fig. S2B).

**Fig. 2.**
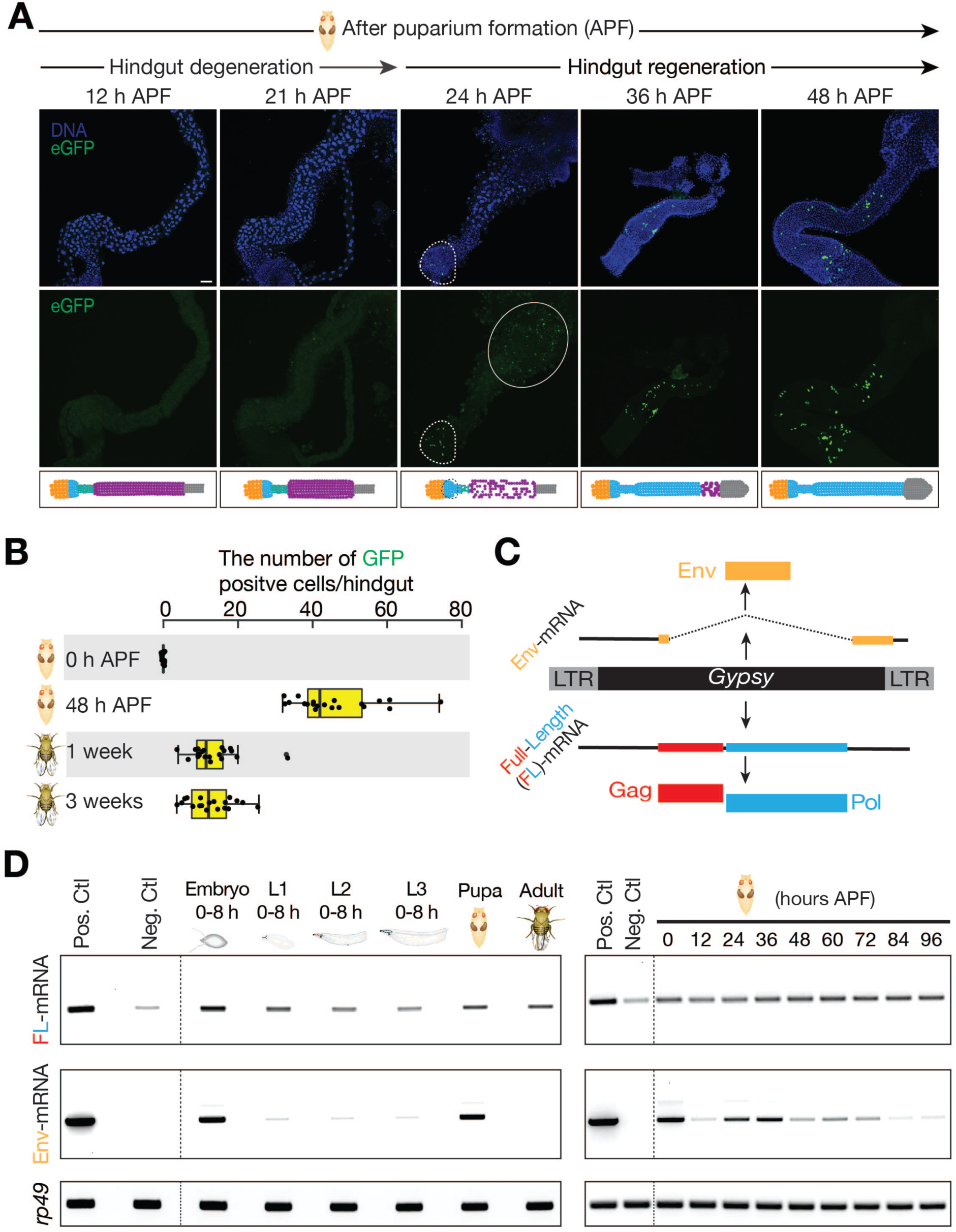
*Gypsy* selectively mobilizes in the regenerating tissues during metamorphosis. **(A)** Monitoring *Gypsy* mobilization in *Drosophila* during hindgut development at the pupal stage via eGFP transposition reporter. No eGFP is expressed during hindgut degeneration (12h and 21h AFP). eGFP positive cells can be detected in the newly formed hindgut (24h, 36h and 48h APF). Dashed circle highlights eGFP signals. solid circle depicts autofluorescene from the dying cells. **(B)** Quantification of the number of eGFP positive cells from *Gypsy* transposition reporter at different stages. **(C)** Diagram depicts the transcripts and proteins from *Gypsy*. **(D)** RT-PCR experiments to measure the expression of full-length and Env-mRNAs from different stages. Pos. Ctl: Positive control, ovaries with Piwi being depleted in follicle cells; Neg. Ctl: negative control, ovaries with *Gypsy* being depleted in both germ cells abd follicle cells. APF, after puparium formation.

The eGFP reporter we employed cannot temporally pinpoint mobilization events and is incapable of determining when mobilization ceases. As adult *Drosophila* hindguts possess no active stem cells and little cell turnover (23), an unceasing flow of *Gypsy* mobilization is predicted to steadily increase the number of eGFP-positive cells as pupae develop into adulthood. However, we detected that the average number of eGFP-positive hindgut cells decreased from 46 in pupa to 13 in adult (Fig. 2B). As with hindgut, we also observed more eGFP-positive cells in pupal salivary glands, proventriculi, and midguts than the corresponding adult structures (Fig. S2C). Therefore, our data indicate that a proportion of cells harboring *Gypsy* mobilization events are eliminated during development, and that *Gypsy* only mobilizes at pupal stage.

To further investigate the activity of *Gypsy* during development, we monitored its mRNA levels at different stages (Fig. 2C and 2D). In addition to transcribing full-length transcripts that encode Gag and Pol proteins, for its activation, *Gypsy* needs to produce spliced transcripts (Env-mRNAs) encoding Env proteins (Fig. 2C) (24). We found that full-length *Gypsy* mRNAs were transcribed throughout all life stages (Fig. 2D), arguing against transcriptional silencing. However, besides being maternally deposited into embryos (data not shown), the Env-mRNAs were primarily produced at pupal stage (Fig. 2D). During *Drosophila* development, the metamorphosis process is driven by a pulse of the steroid hormone (ecdysone) at 0 hours and 24 hours APF (25). Interestingly, we found that temporal waves of Env-mRNA expression correlated with ecdysone peaks (Fig. 2D).

What is the function of *Gypsy* activation during metamorphosis? To get an answer, we embarked on experiments in which we use RNAi constructs to ubiquitously suppress *Gypsy*. Being able to deplete over 80% of *Gypsy* mRNAs and completely abolish *Gypsy* mobilization during development (Fig. 3A and Fig. S3A), we concluded that the RNAi construct we designed could efficiently silence *Gypsy*. With *Gypsy* being suppressed (sh-*Gypsy*), we found that flies still developed normally and showed no overt phenotypes.

**Fig. 3.**
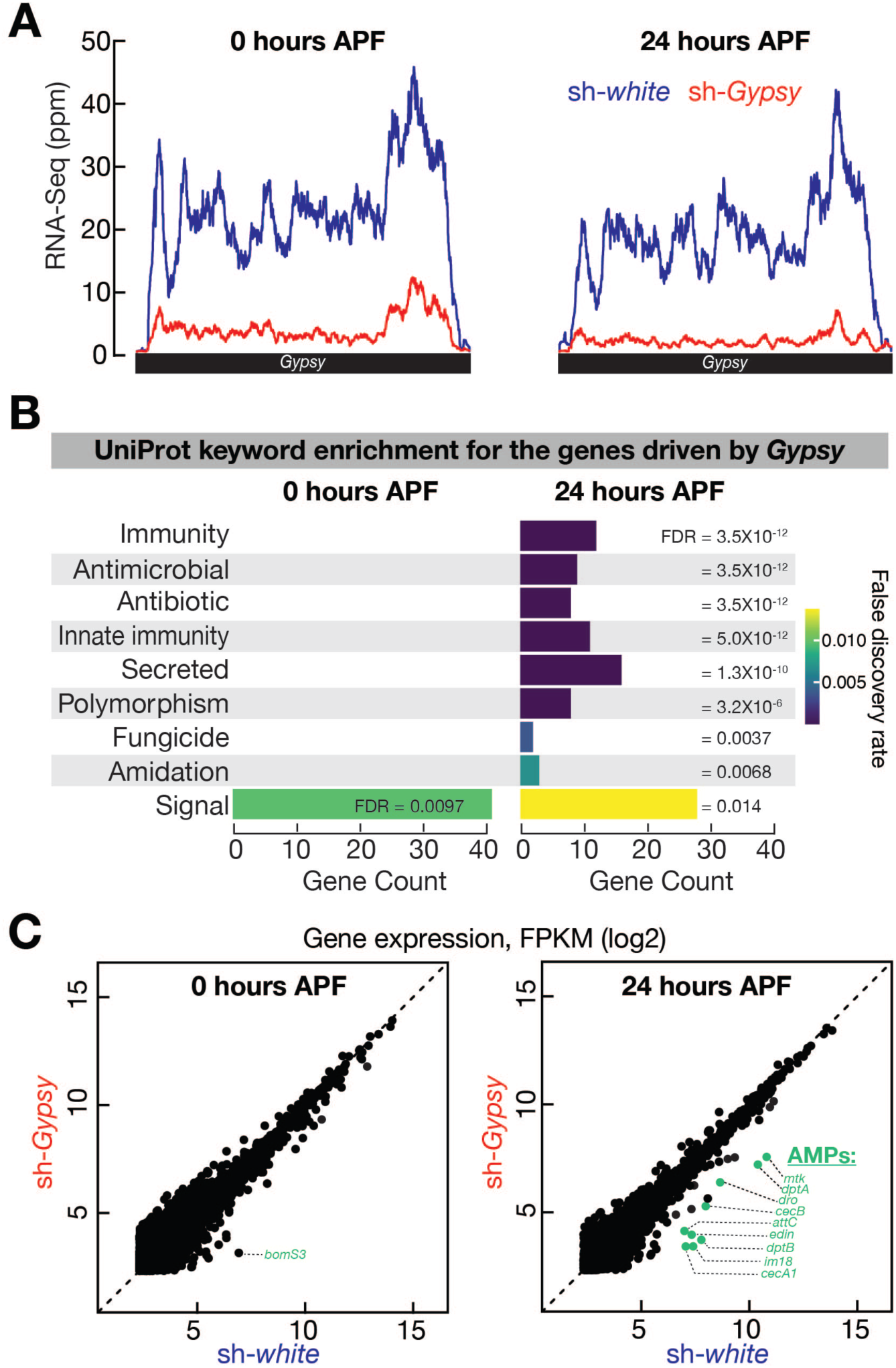
Activation of *Gypsy* during metamorphosis ptimates inate immunity. **(A)** RNA-seq profiles of *Gypsy* expression from 0 and 24 hours of *white* and *Gypsy* depleted pupae. sh-RNAs are driven by *ac*-Gal4. **(B)** The analysis of UnitProt keyword enrichment of differentially expresed genes (more than 2-fold change) between *white* and *Gypsy* depleted pupae. **(C)** Transcriptome profiles from *white* and *Gypsy* depleted pupae. Genes with FPKM greater than 5 are plotted. AMPs: antimicrobial peptides. The genes decreased for more than 5-fold in *Gypsy* depleted pupae are shown in green color.

To examine potential changes of the host transcriptome upon *Gypsy* suppression, we performed RNA-Seq experiments using flies with the *white* being silenced (sh-*white*) as controls. To determine the potential function of *Gypsy* activation at pupal stage, we performed functional enrichment analysis on the genes whose expression is promoted by *Gypsy* (Fig. 3B). At 0 hours APF, the only enriched term from the UniProtKB Keyword classification system was “signal” (FDR = 0.01, Fig. 3B). Strikingly, at 24 hours APF, multiple terms relating to immunity were significantly enriched among the list of genes whose expression decreased upon *Gypsy* depletion: immunity, antimicrobial, antibiotic, and innate immunity (all with FDR < 5×10^−12^, Fig. 3B). Detailed examination revealed that *Gypsy* activation in fly pupae drives expression of the antimicrobial peptides (AMPs, Fig. 3C and Fig. S3B), the effector proteins of innate immunity to combat invading pathogens. (26–28). At 0 hours APF, among the 6 genes that were differentially expressed for more than 5-fold, only 1 of them was AMP (Fig. 3C). However, at 24 hours APF, we found that among the 11 genes that were significantly decreased more than 5-fold upon *Gypsy* suppression, 9 of them (82%) were AMPs (Fig. 3C and Fig. S3B). Based on these findings, we conclude that *Gypsy* activation during metamorphosis triggers a wave of innate immune response.

Recently it was proposed that in both flies and mice, activation of innate immunity during a single event could protect animals from future pathogen infection (29, 30). Accordingly, we hypothesized that *Gypsy* activation during metamorphosis would enable adult flies to rapidly mount an immune response to combat pathogens. We tested this hypothesis by infecting 5-day-old adult flies with one dose of *<u>D</u>rosophila* <u>C V</u>iruses (DCV, Fig. 4A), a group of RNA virus that is pathogenic to fruit flies (31). Control flies (sh-*white*) that experienced normal *Gypsy* activation at the pupal stage cleared the viruses within one day (Fig. 4B). In contrast, flies in which *Gypsy* was silenced were unable to clear viruses (Fig. 4B). Based on RT-qPCR experiments, at 3 days after infection, the ∆∆Ct value for DCV RNA levels between sh-*Gypsy* and sh-*white* flies is 20.27 (equivalent to an 1,265,829-fold difference. sh-*Gypsy*: DCV ∆Ct = −4.95; sh-*white*: DCV ∆Ct = 15.32, Fig 4C). At 6 days after infection, there was 1,878,976-fold more DCV in sh-*Gypsy* flies than controls (sh-*Gypsy*: DCV ∆Ct = −5.86; sh-*white*: DCV ∆Ct = 14.98; and ∆∆Ct = 20.84, Fig 4C). At 10 days after infection, there was a 62,557-fold difference (sh-*Gypsy*: DCV ∆Ct = – 3.77; sh-*white*: DCV ∆Ct = 12.17; and ∆∆Ct = 15.94, Fig 4C). During these first 10 days, sh-*Gypsy* flies showed significantly 4.4-fold higher mortality rate than sh-*white* controls: 28.8% for sh-*Gypsy* flies vs. 6.6% for sh-*white* animals (*p* = 4.3×10^−7^, Cox Proportional analysis; Fig. S4A and S4B). Subsequently, the mortality rates of sh-*Gypsy* flies were decreased to a level that is indistinguishable from sh-*white* control animals (Fig. S4A and S4B). Consistently, there was minimal, if any, DCV detectable in the sh-*Gypsy* survivors at 15 days after infection (Fig. 4B and 4C), indicating that the survivors eventually clear out DCV. In summary, our findings indicate that *Gypsy* activation at pupal stage renders a robust ability to rapidly defeat invading pathogens at adult stage.

**Fig. 4.**
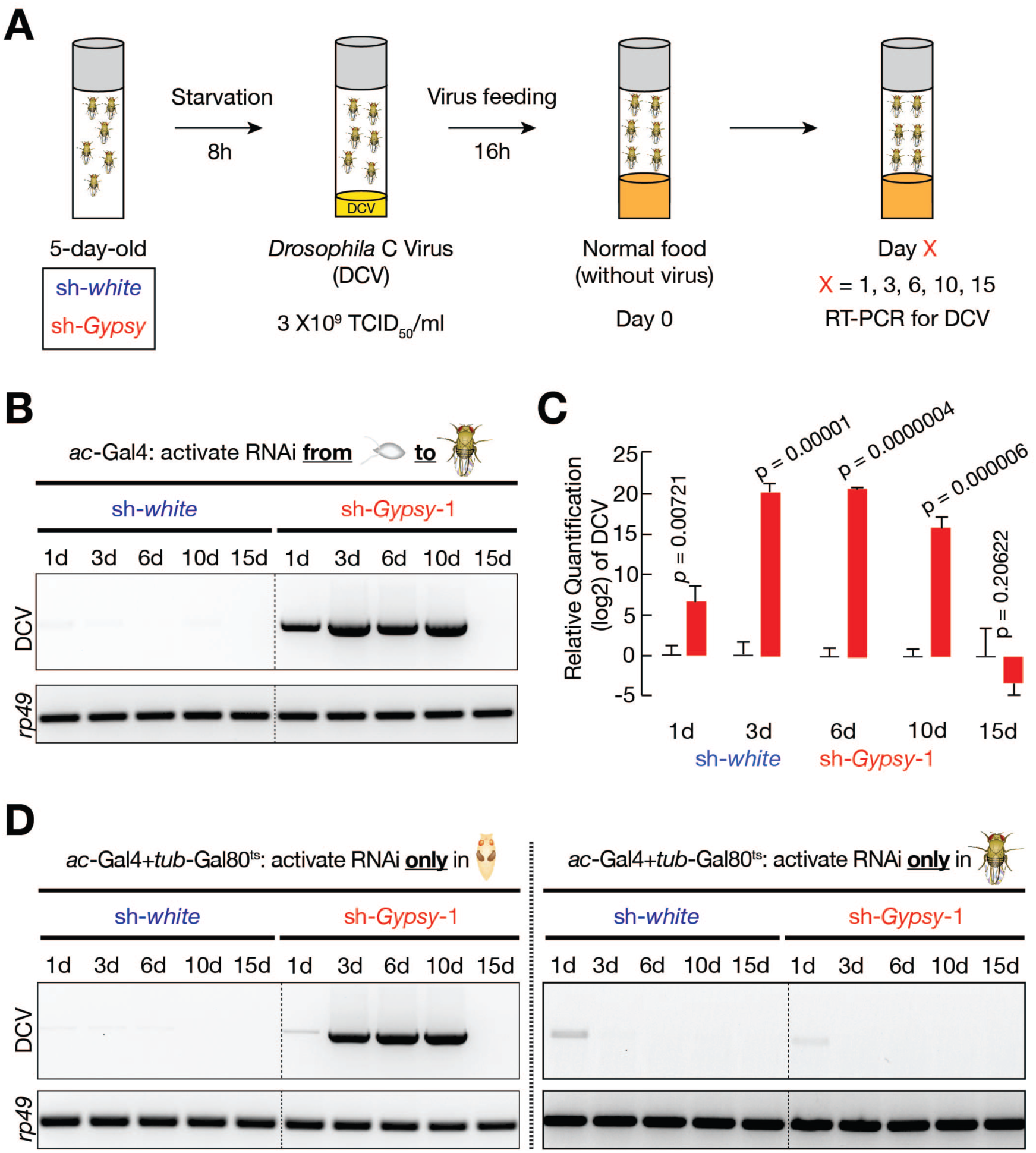
*Gypsy* activation at pupal stage contributes to viral clearance in adult flies. **(A)** Experimental scheme. **(B)** RT-PCR to examine the DCV mRNA levels in the fly body, rp49 mRNA levels serve as loading control. The RNAi constructs are active during the whole *Drosophila* life cycle. Survivors cleared out the virus by day 15. **(C)** RT-qPCR to quantify the fold changes of DCV mRNA in sh-*Gypsy* flies, relative to sh-*white* controls. **(D)** RT-PCR to examine the DCV mRNA levels in the fly body upon suppressing *Gypsy* at either pupal (left panel) or adult (right panel) Stage.

Besides just one dose of viral infection, we also tested the condition of chronic infection by housing adult flies on virus-containing food for 20 days (Fig. S4C). Under this circumstance, *Gypsy* activation at pupal stage was also essential to optimally protect the hosts. For the control group (sh-*white*), only 30% of animals died at the end of 20 days infection (Fig. S4C). However, for flies with *Gypsy* being silenced (sh-*Gypsy*) at pupal stage, the lethality rate significantly soared to 78% (*p* = 7.2×10^−9^, Cox Proportional analysis; Fig. S4C).

The aforementioned experiments on testing the function of *Gypsy* were performed by using RNAi strategy to suppress *Gypsy* expression. RNAi technology potentially has off-target effects as a shortcoming, and this can be addressed by employing multiple RNAi constructs that target different regions of the same gene. Therefore, we further designed 6 other RNAi constructs to silence *Gypsy* during animal development and orally infected adult flies with one dose of DCV. Consistently, we found that decreasing *Gypsy* activity compromises the antiviral response in adult flies (Fig. S5). Interestingly, it appears that the RNAi efficiency negatively correlates with the robustness of antiviral response. For example, for the two RNAi constructs (sh-*Gypsy*-4 and sh-*Gypsy*-7) that only deplete *Gypsy* mRNA less than 50%, flies had less DCV accumulated in their body (Fig. S5). These data further support our conclusion that *Gypsy* activation is essential for mounting a robust antiviral response.

Although our data show that *Gypsy* specifically produces Env-mRNAs and mobilizes at pupal stage (Fig. 2), its full-length transcripts are constantly expressed throughout the *Drosophila* life cycle (Fig. 2D). Thus, we hypothesized that these transcripts or the proteins encoded by them acting in adults, but not *Gypsy* activation at pupal stage, contributed to DCV clearance. To test this idea, we performed experiments to suppress *Gypsy* activity in a stage-specific manner (Fig. S6, S7A, and S7B). When silencing *Gypsy* during metamorphosis, but allowing full-length transcripts expression in adulthood, flies lacked the ability to clear the orally infected DCV (Fig. 4D and Fig. S7C). On the other hand, flies with full-length transcripts suppressed at adulthood, but had experienced *Gypsy* activation at pupal stage, had no defects on DCV clearance (Fig. 4D and Fig. S7C). We conclude that it is the *Gypsy* activity during metamorphosis, but not other stages, that bestows upon hosts a robust ability to combat invading viruses.

The current view posits that transposons are largely suppressed during development to minimize their detrimental effects to the hosts. Our findings suggest a new model for the function of developmental retrotransposon activation. We propose that, during metamorphosis, hosts use a wave of retrotransposon activation to prime their innate immunity in the newly regenerated tissue that will last to adult stages and even entire lifetimes. Similar to the role of vaccination, the immune priming function of retrotransposons provides long-term protection from pathogen infections.

There are several possible mechanisms by which *Gypsy* might trigger an immune response. Given the restricted expression window of Env proteins, they may directly serve as ligands to activate immune signaling. Interestingly, the Env proteins from vesicular stomatitis virus indeed could trigger Toll-7 activation in *Drosophila* (32, 33), supporting this hypothesis. Alternatively, similar to how HIV DNA triggers cGAS/STING-mediated immune response in human cells (34), *Gypsy* DNA may activate dmSTING to drive AMP production. How could *Gypsy*-primed immunity grant adult animals protection upon viral infection? Given that mounting a prompt antiviral response is essential for viral clearance, priming of the immune system by *Gypsy* may launch the response in a timelier fashion upon infection. Besides promoting antiviral-signaling for infected cells, *Gypsy* activation might stimulate the activity of macrophages– the cells that engulf viruses and virus-infected cells. Supporting our hypothesis, evidence indicates that macrophages from fly embryos can engulf microbes only if they have previously been “primed” by components from dead cells (30).

During human embryogenesis, there is also a wave of *Gypsy*-like retrotransposon activation–from 8-cell stage to blastocyst stage, at which time the paternal genome is activated (3). As maternal immunity is suppressed at these stages to avoid immune attack, it is possible or likely that developing mammalian embryos employ an analogous mechanism to *Drosophila* to gain a long-term protection from pathogen infection. Although detailed mechanisms may differ, harnessing transposons to prime immune systems for subsequent pathogen warfare is likely a recurrent theme from flies to mammals.

---

**Fig. S1.**
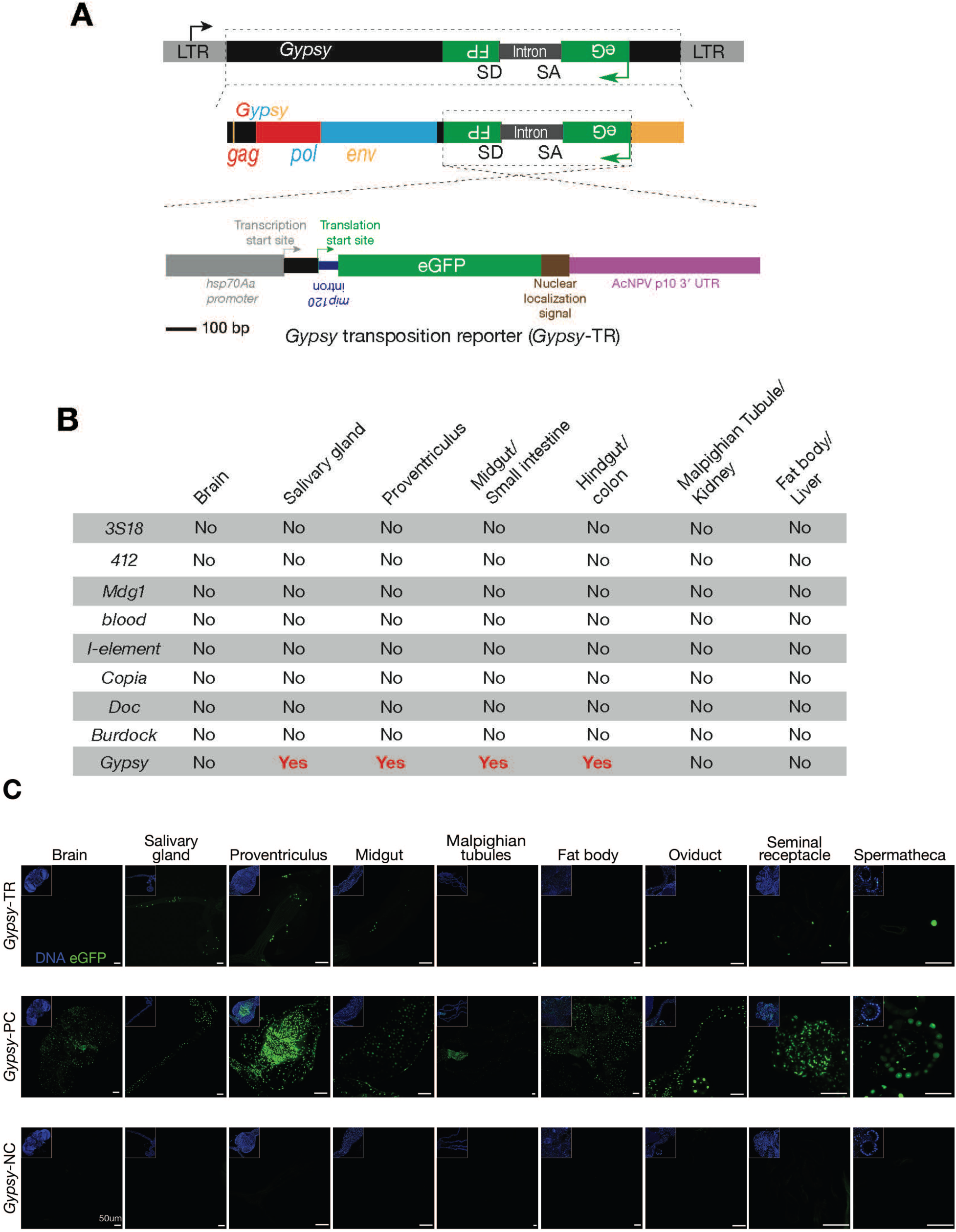
Monitoring retrotransposon mobilization in somatic cells via a transposition reporter. **(A)** Detailed schematic design of eGFP transposition reporter to monitor *Gypsy* mobilization. **(B)** Summary of mobilization events from different somatic tissues for 9 retransposon families, as assayed by corresponding eGFP reporter. No: no eGFP positive cells are detected; Yes: eGFP positive cells can be detected. **(C)** Detecting eGFP signals in somatic tissues from positive control, negative control, and *Gypsy* transposition reporterin 2-4 day adult flies. Note:Positive control constructs gives low number of eGFP positive cells in brain and malpighian tubules, indicating the transcription of *Gypsy* is suppressed in these tissues.

**Fig. S2.**
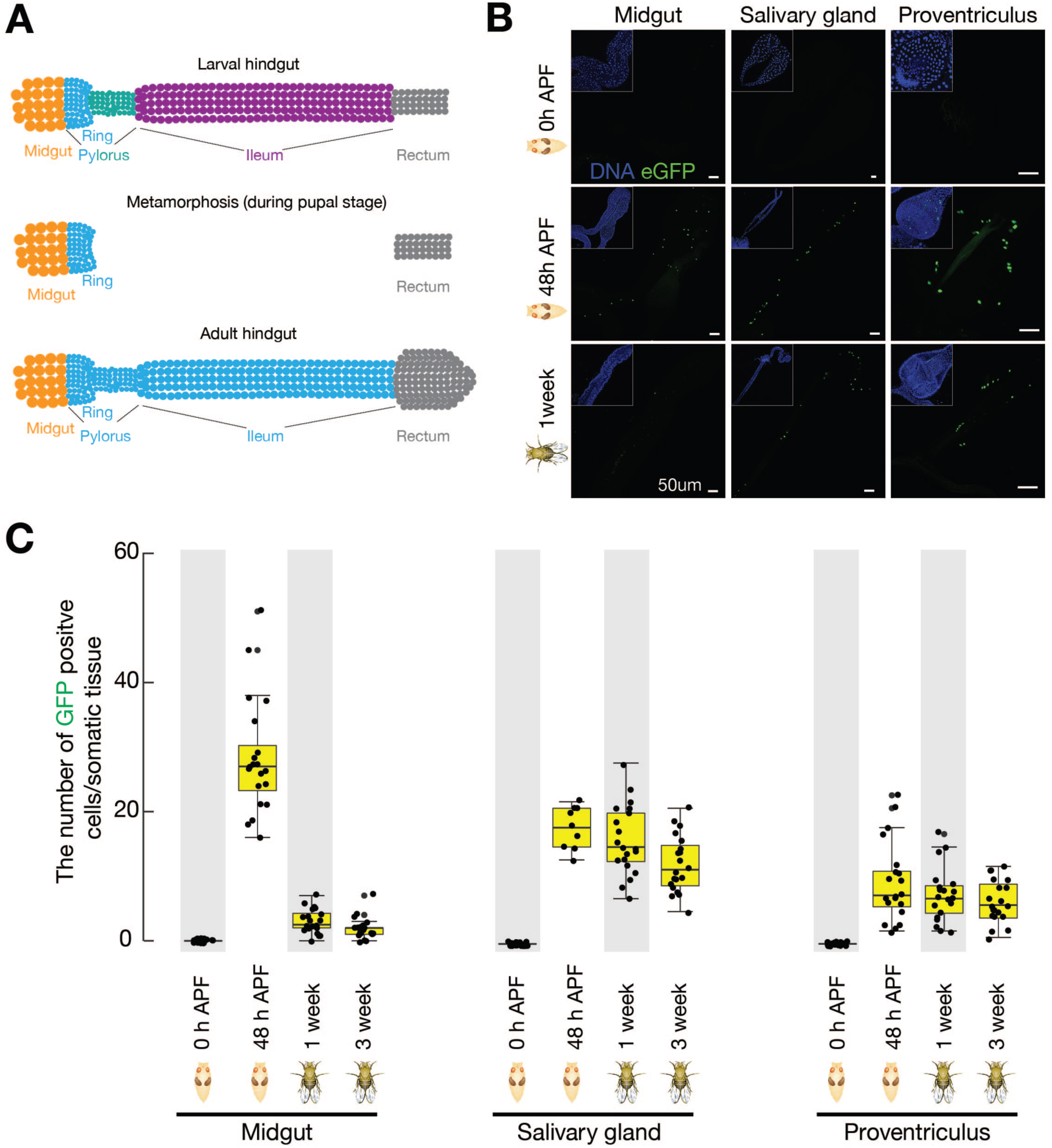
*Gypsy* selectively mobilizes in the regenerating tissues during metamorphosis. **(A)** Schematic of *Drosophilla* hindgut. Both larval and adult hindgut include the pylorus, ileum and rectum. During pupal stage metamorphosis, the pylorus and ileum from larval stagedegenerate; the anterior part of pylorus (ring) regenerates to produce adult pylorus and ileum. **(B)** Detecting eGFP positive cells from *Gypsy* transposition reporter in midgut, salivary gland and proventriculus at deifferent stages. **(C)** Quantifying the number of eGFP positive cells from *Gypsy* transposition reporter in midgut, salivary gland and proventriculus.

**Fig. S3.**
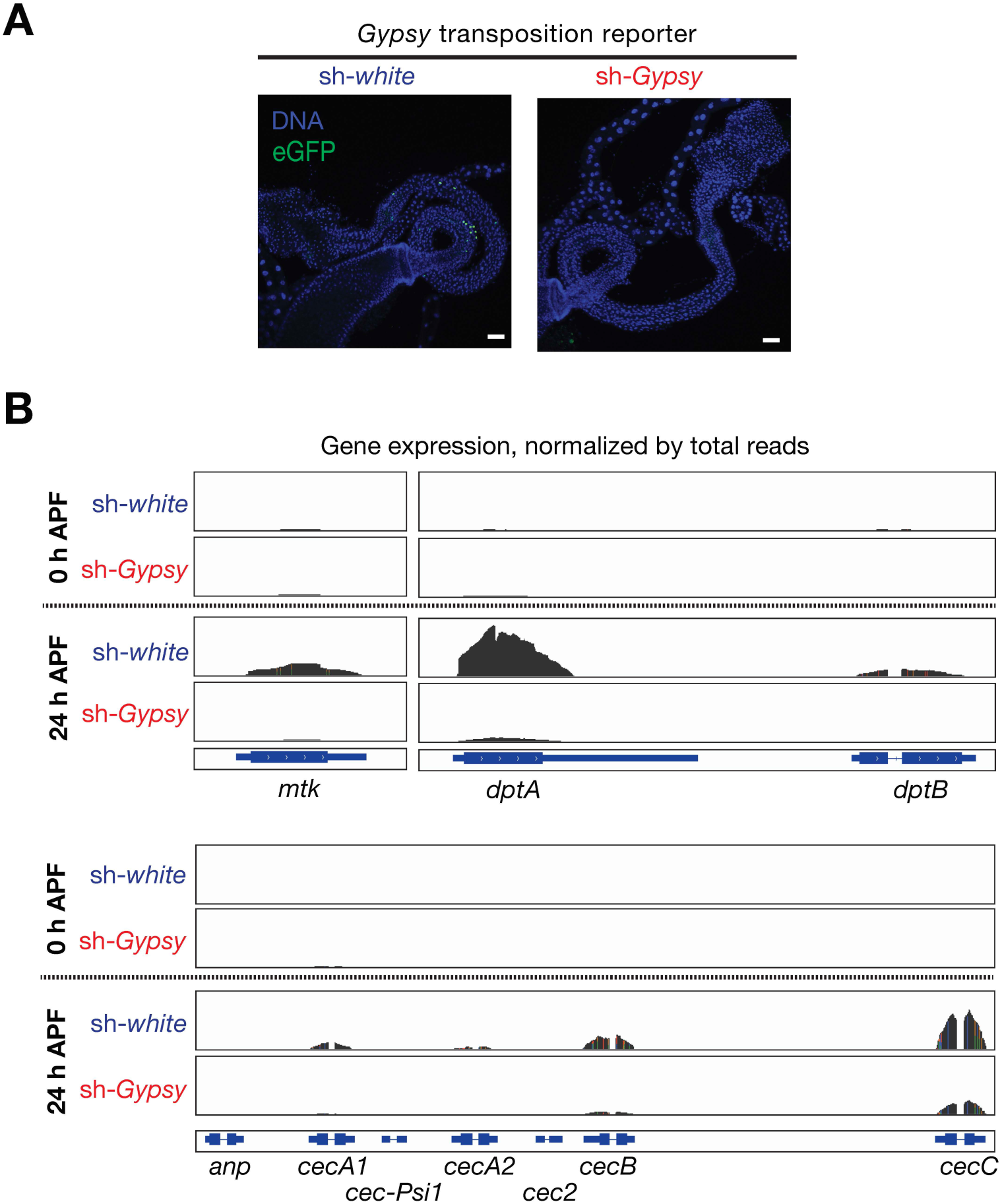
*Gypsy* activation is required for driving AMP production in fly pupae. **(A)** Visualizing *Gypsy* transposition events (GFP postive cells) in hindgut fron either *white* or *Gypsy* suppressed2-4-day old adults. **(B)** RNA-Seq profiles to examine the expression of AMP genes. APF, after puparium formation.

**Fig. S4.**
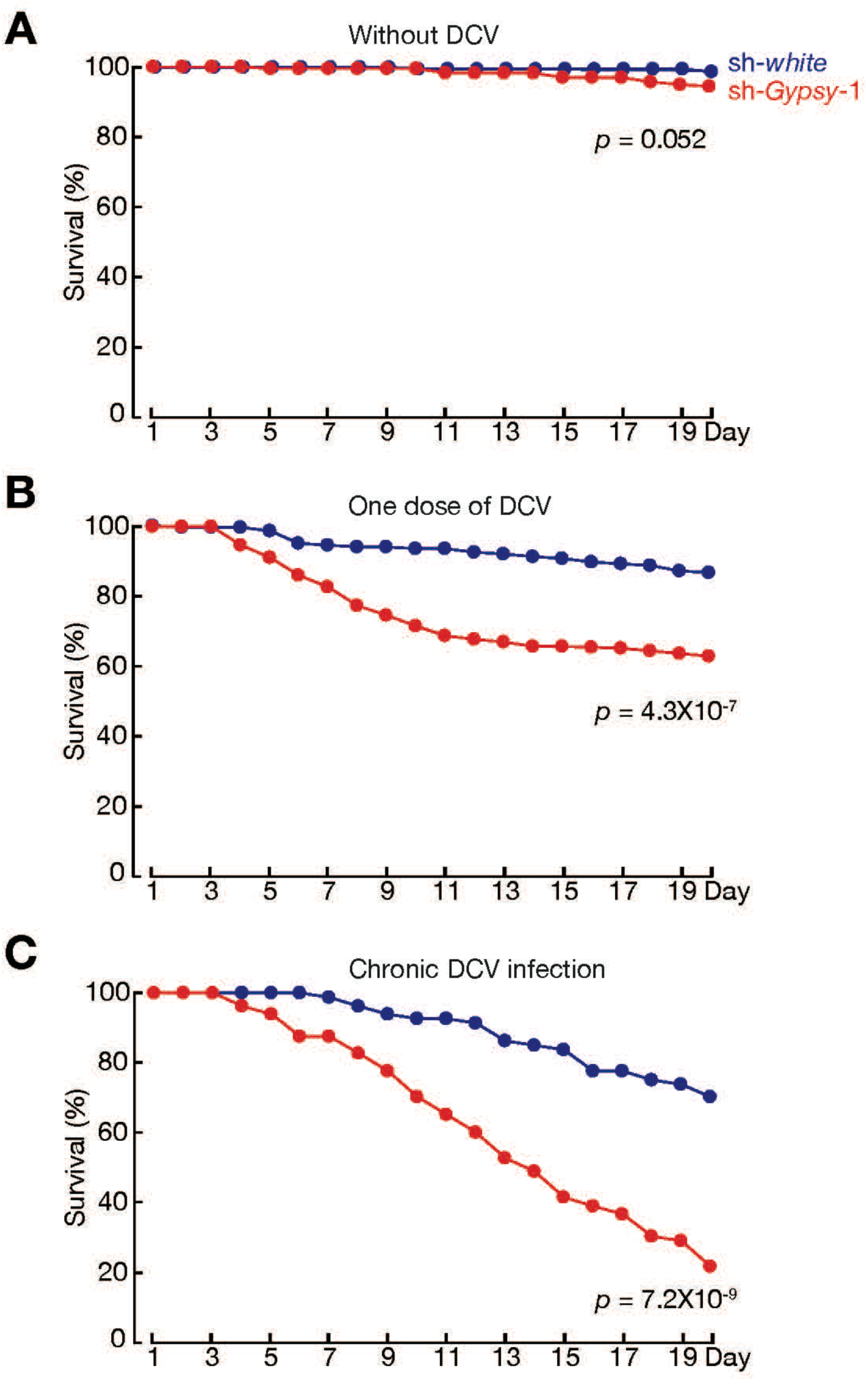
The survival rates of white and Gypsy depleted flies at three DCV oral infections conditions. **(A)** Starvation but without DCV infection. **(B)** Starvation and with one dose of DCV infection. **(C)** No starvation and with chronic infectionby raising flies on DCV-containing food for 20 days.

**Fig. S5.**
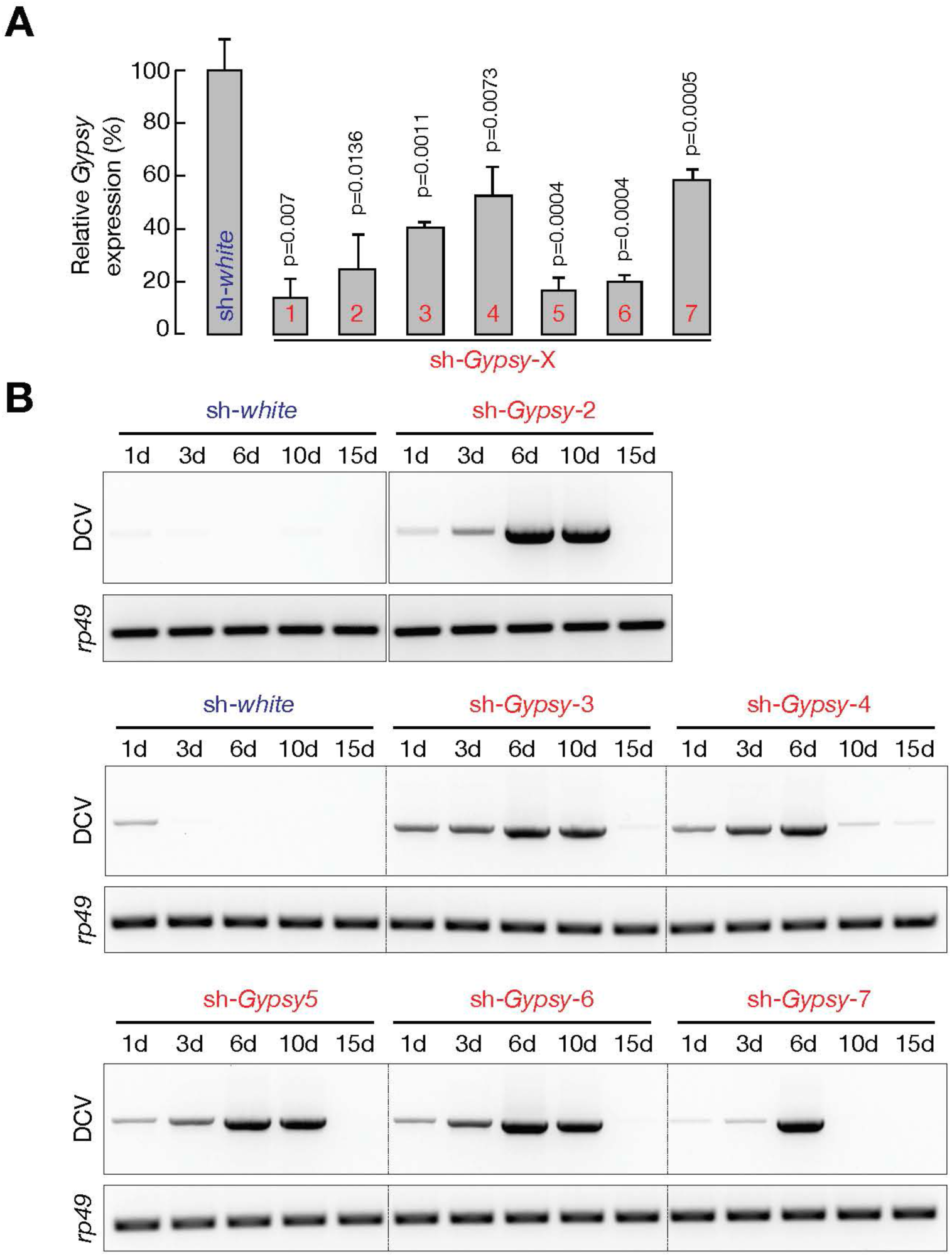
The immune-priming function of *Gypsy* contributes to viral clearance in adult flies. **(A)** RT-qPCR to quantify the expression of *Gypsy* upon suppression by using one of the 7 RNAi constructs. **(B)** RT_PCR to examine DCV mRNA levels.

**Fig. S6.**
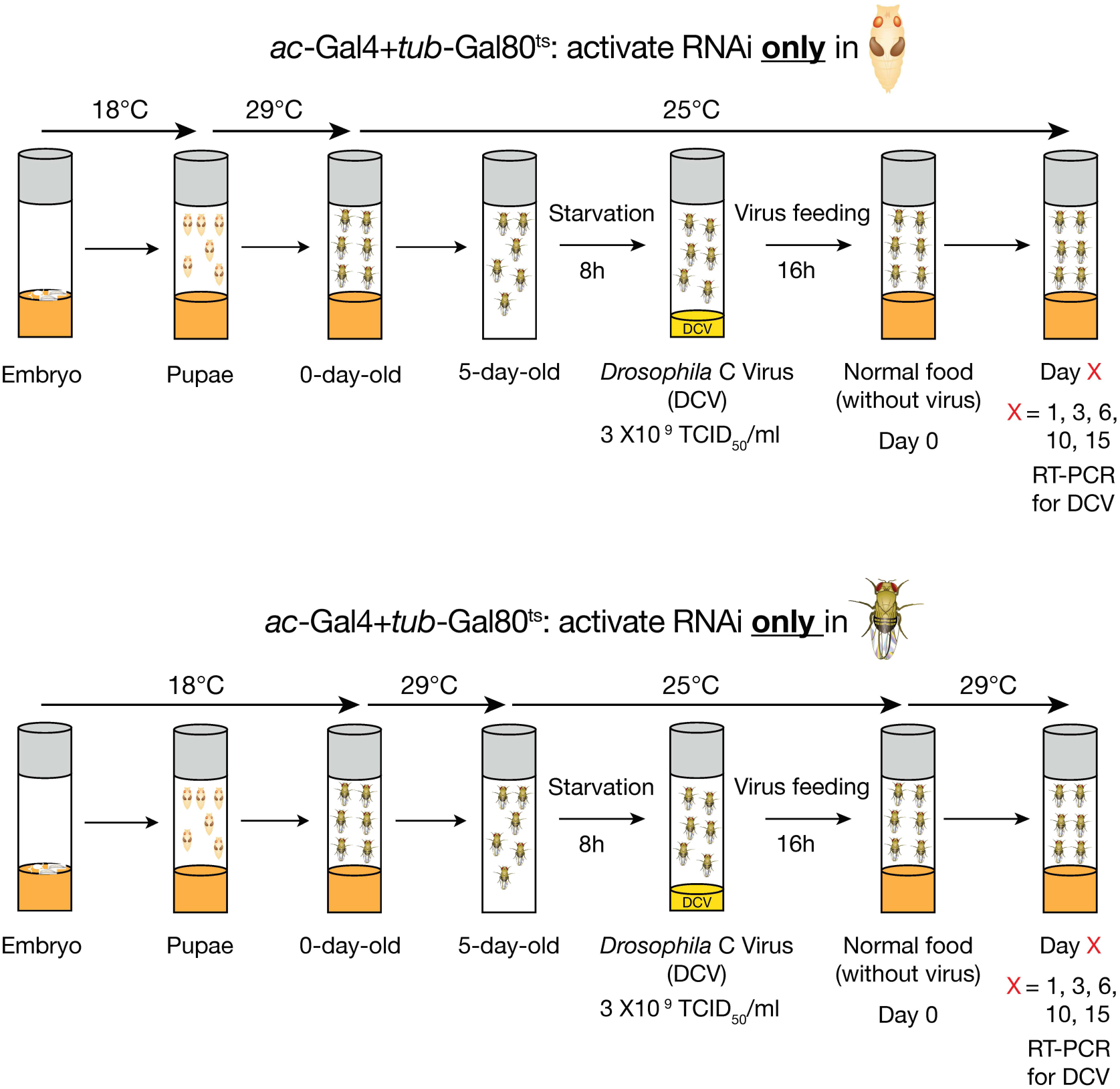
Schematic design of stage-specific depletion protocol by using Gal4-Gal80^ts^ system. At low temperature (18°C), Gal80 inhibits Gal4 activity. At 29°C, Gal80 becomes inactive and cannot suppress Gal4.

**Fig. S7.**
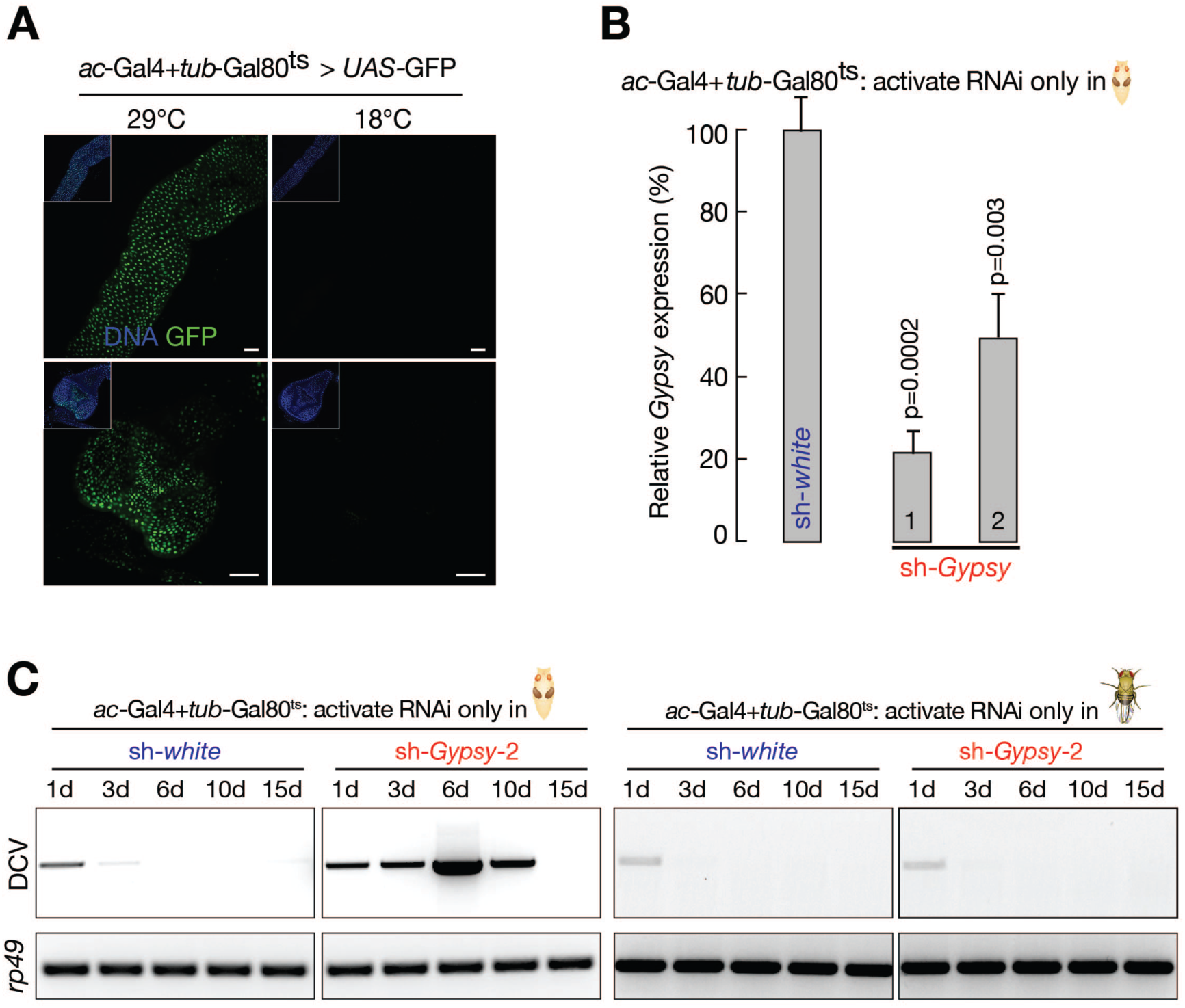
The immune-priming function of *Gypsy* contributes to viral clearance in adult flies. **(A)** Validating Gal4-Gal80^ts^ system by driving *uasp*-GFP expression at high temperature (29°C) and low temperature (18°C). **(B)** Quantification of *Gypsy* expression from flies with *white* and *Gyspy* being specifically depleted at pupal stage by using RT-qPCR. **(C)** RT-PCR to examine the DCV mRNA levels in the fly body upon stage-specifically suppressing *Gypsy*.

## Acknowledgments

We thank Phillip Zamore for providing DCV. We thank members from ZZ lab, Don Fox, and Neal Silverman for critical suggestions, Brandy Kegeris for assistance on cloning, and Ken Poss for reading manuscript. This work was supported by the grant from the National Institutes of Health to Z.Z. (DP5 OD021355).

## Author Contributions

Z.Z. and L.W. conceived the project and designed the experiments. L.T. performed one replicate of RNA-Seq experiments for 24 hours pupae, and helped with data analysis for Fig. 3B. L.W. performed all of the rest experiments and analyzed data. Z.Z. and L.W. wrote the manuscript.

